# UNC-45 has a crucial role in maintaining muscle sarcomeres during aging in *Caenorhabditis elegans*

**DOI:** 10.1101/2022.06.04.494828

**Authors:** Courtney J. Matheny, Hiroshi Qadota, Marion Kimelman, Aaron O. Bailey, Andres F. Oberhauser, Guy M. Benian

## Abstract

As people live longer, age-related diseases, like sarcopenia, will become a greater public health concern. We use the model organism *C. elegans* to better understand the molecular mechanisms behind muscle maintenance. Muscle function is dependent on having properly organized and functioning thick filaments, which are primarily composed of myosin. The myosin head requires the chaperone UNC-45 to initially fold it after translation and is likely used to re-fold back to functionality after thermal or chemical stress induced unfolding. We observe an early onset of sarcopeania when UNC-45 is perturbed during adulthood. We observe that during adult aging, there is a sequential decline of HSP-90, UNC-45, and then myosin. Myosin and UNC-45 protein decline are independent of steady state mRNA levels. Loss of UNC-45 is correlated with an increase in phosphorylation of the protein. By mass spectrometry, S111 was identified as being phosphorylated and this modification may affect binding to HSP-90. A longevity mutant with delayed onset of sarcopenia also shows a delay in the loss of HSP-90, UNC-45, and myosin. We also see a decrease in UNC-45 protein, but not transcript, in an *hsp-90* loss of function mutant, suggesting a role for HSP-90 in stabilizing UNC-45. This leads us to propose the model that during aging, a loss of HSP-90 leads to UNC-45 being post translationally modified, such as phosphorylation, and degraded, which then leads to a loss of myosin, and thus muscle mass and function. A better understanding of how myosin and its chaperone proteins are regulated and affected by aging will lead to better preventative care and treatment of sarcopenia and, possibly, the age-related decline of heart muscle function.

**Graphical Abstract:** 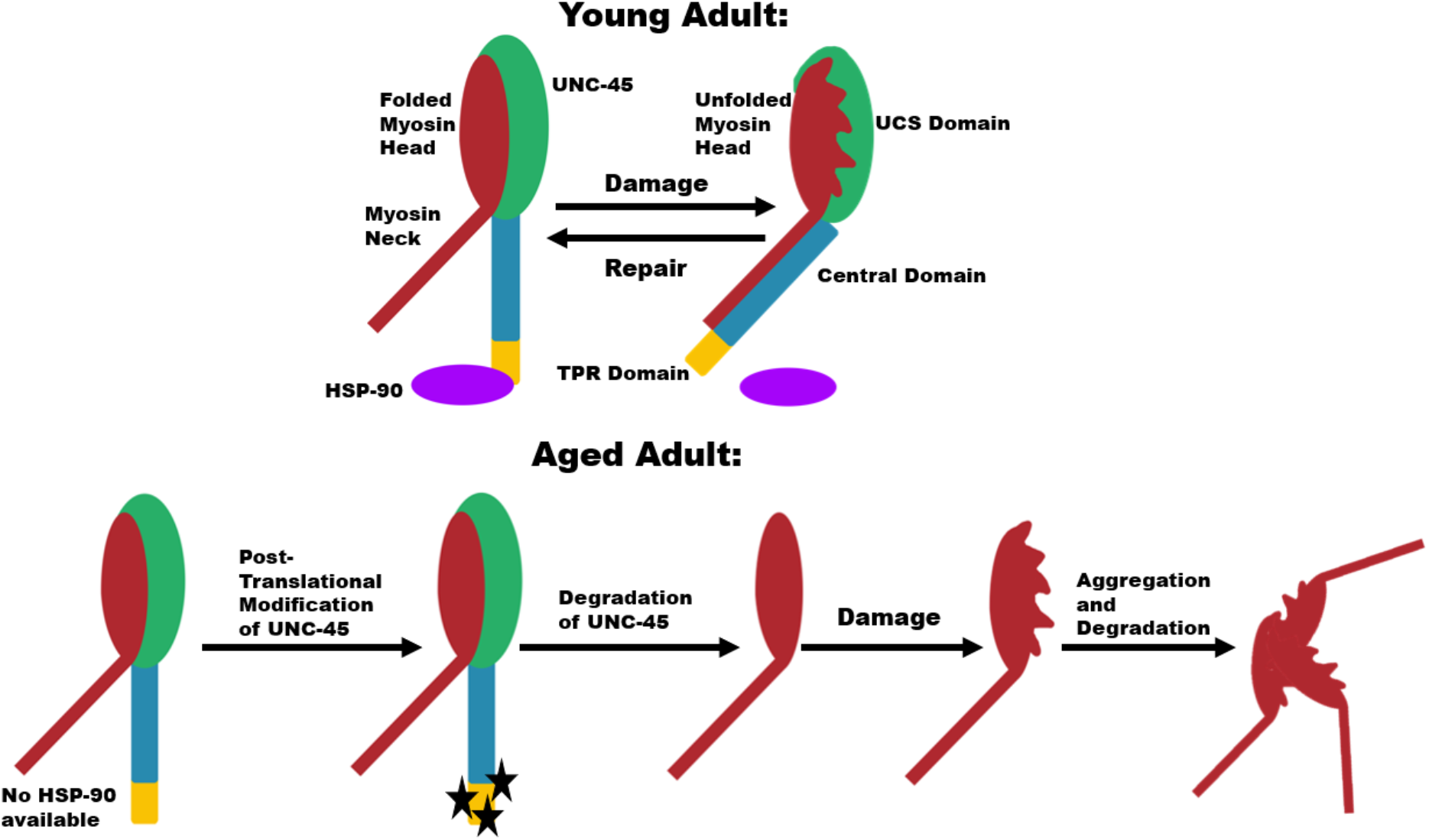

In young adults, under normal conditions the UCS domain of UNC-45 (shown in green) is bound to the myosin head (in red) and the TPR domain (in yellow) is bound to HSP-90 (in purple). Under stress conditions, HSP-90 detaches from the TPR domain, causing a conformational change in UNC-45 that allows the Central domain (in blue) to bind to the myosin neck (in red) resulting in inhibition of the myosin power stroke while the UCS domain protects/re-folds the myosin head. HSP-90 can then rebind the TPR domain, causing the Central domain to release the myosin neck, allowing movement of the myosin motor. However, aged adults experience a loss of HSP-90 and UNC-45 (which has increased post translational modification with aging). The loss of the Myosin chaperones leads to increased aggregation and degradation of Myosin with age.This loss of Myosin at the thick filament results in decline in muscle mass and function, also known as sarcopenia. Note that only the myosin head and neck are shown for simplicity of illustration.

## Introduction

Sarcopenia, the decline in skeletal muscle mass and function without any underlying disease, is a major contributor to physical disability, poor quality of life, and death among the elderly (Cruz-Jentoft et al., 2014). 40-50% of individuals over 80 years of age suffer from this loss of muscle mass and function (Barbosa-Silva et al., 2016; Iannuzzi-Sucich et al., 2002). The molecular mechanisms responsible for this age-related condition remain uncertain (Zembron-Lacny et al., 2014). Resistance training and dietary changes are recognized as the gold standard therapy but have only a modest effect (Candow, 2011). There is a direct association between poor hand grip strength, reduced physical function and a higher risk of falling (Szulc et al., 2016). Intriguingly, even in middle age (40-69), there is a correlation between reduced grip strength and all-cause mortality and incidence of and mortality from cardiovascular disease, respiratory disease, and cancer (Celis-Morales et al., 2018). Elderly individuals at a higher risk of falling are at a higher risk of vertebral and non-vertebral fractures (Szulc et al., 2016) – leading to surgeries, hospitalization, and increased medical complications and risks. Additionally, the increased risk of respiratory illness in individuals over 65 years of age may be partially explained by the ageing-related weakening of the diaphragm muscle resulting in non-productive coughs and more severe respiratory illnesses (Gosselin et al., 1994). With the ever-increasing population of elderly and the predicted strain on the healthcare system (Statistics, 2012), it is crucial we understand the molecular mechanisms responsible for age-related diseases like sarcopenia so that we can develop more effective therapies and prevention methods.

Sarcopenic patients exhibit a loss of myofibrils, which are comprised of thin and thick filaments. The thin filaments are predominantly composed of filamentous actin while thick filaments are predominantly composed of myosin (Henderson et al., 2017). There is a superfamily of myosins composed of at least 30 classes of myosins, but all myosins consist of 3 regions: a “head”, a “neck”, and a “tail” (Squire et al., 2017). The head is a complex structure that converts the energy of ATP hydrolysis to the mechanical work of binding and moving along F-actin tracks (Squire et al., 2017). The chaperone UNC-45 is required to fold the myosin head initially after translation and, likely, to re-fold the myosin head after stress results in unfolding (Barral et al., 1998; Barral et al., 2002; Etard et al., 2008; Kachur and Pilgrim, 2008). UNC-45 may also be crucial in mature muscle because of the physical stress muscle cells undergo throughout life and the relatively slow turnover rate of myosin in established thick filaments (Solomon and Goldberg, 1996). UNC-45, which is conserved across all eukaryotes, was first identified in *Caenorhabditis elegans* and named for the uncoordinated (impaired movement) phenotype observed in mutant animals (Epstein and Thomson, 1974). Mis-regulation of UNC-45 is associated with several diseases including myopathies, cardiomyopathies, cataracts, and cancer metastasis (Bazzaro et al., 2007; Bernick et al., 2010; Esteve et al., 2018; Hansen et al., 2014; Janiesch et al., 2007; Melkani et al., 2011; Wohlgemuth et al., 2007). UNC-45 is comprised of a C-terminal UCS domain responsible for binding myosin, an N-terminal tetratricopeptide repeat (TPR) domain that interacts with the heat shock co-chaperone protein HSP-90 (Barral et al., 2002), and a central domain that acts as an inhibitor of the myosin power stroke (Bujalowski et al., 2018). These authors put forth the following model: Under normal conditions, the UCS domain of UNC-45 is bound to the myosin head and the TPR domain is bound to HSP-90. Under stress conditions, HSP-90 detaches from the TPR domain, causing a conformational change in UNC-45 that allows the Central domain to bind to the myosin neck resulting in inhibition of the myosin power stroke while the UCS domain re-folds the myosin head. After refolding of the myosin head, HSP-90 then rebinds the TPR domain, causing the Central domain to release the myosin neck, allowing movement of the myosin motor (Bujalowski et al., 2018).

*C. elegans* is an excellent genetic model organism to study sarcomere assembly, maintenance and regulation (Gieseler et al., 2017), and conserved mechanisms of aging (Kenyon, 2010). The *C. elegans* model provides the shortest lifespan and largest possible sample size among the models used to study sarcopenia. Their short lifespan (average 18-21 days) makes them particularly convenient for aging studies. Muscle function is easy to monitor in worms since they require functioning body wall muscles for locomotion. Additionally, nematode muscle does not contain stem cells and thus provides an opportunity to investigate how the assembled muscle contractile apparatus is maintained and functions during aging in the absence of regeneration. Monica Driscoll’s lab was the first to report that *C. elegans* undergo an age-dependent decline in whole animal locomotion and deterioration of the muscle myofilament lattice and thus *C. elegans* is a good model for sarcopenia (Herndon et al., 2002).

Here we have verified and expanded upon evidence that *C. elegans* develop sarcopenia as they age. We observe early onset of sarcopenia when UNC-45 is perturbed at the beginning of adulthood in a temperature sensitive mutant, providing evidence that UNC-45 is important during adulthood and that UNC-45 may play a role in sarcopenia pathology. We have characterized the change in mRNA and protein levels of UNC-45, its co-chaperone HSP-90, and its myosin clients during aging. Additionally, we have begun to determine the mechanism responsible for the decline in UNC-45 during aging.

## Results

### *C. elegans* develop sarcopenia and show decreased numbers of assembled thick filaments during aging

In the studies reported in Herndon et al. (2002), a transgenic line overexpressing GFP-MHC A was used to show that old adults (day 18) had disorganization of thick filaments, as compared to day 4 adults. In retrospect, this result could be questioned because in this strain, even young adults show some disorganization of thick filaments (unpublished data). Therefore, we have verified and extended the results reported in Herndon et al. (2002) by performing immunostaining on wild type worms using an antibody to MHC A (one of the myosin heavy chain isoforms in body wall muscle), and also examined earlier and additional time points. We observe that the number of A-bands, or assembled thick filaments, declines with age (Figure 1A-F). We began counting A-bands once maturity was reached at day 1 of adulthood and continued until the majority of animals reached a sarcopenic state at day 16. A significant loss of A-band number can seen by day 8 of adulthood. Elderly animals also show a decline in thick filament organization at days 12 and 16 (Fig.1 D&E), similar to *unc-45* missense mutants (Landsverk et al., 2007). These results support the notion that *C. elegans* experience a loss of muscle mass with age, beginning at the midpoint of life and continuing until death, analagous to humans. We also measured whole animal locomotion or motility using a crawling assay and found that motility declines by day 4 of adulthood and continues to decline until there is almost no significant mobility by day 16 (Figure 1G). Interestingly, muscle function appears to decline before muscle mass loss is seen. This may suggest that some thick filaments are rendered less functional before they are lost and/or, very likely, that loss of muscle mass is not the sole contributor to loss of muscle function with age.

**Figure 1.**
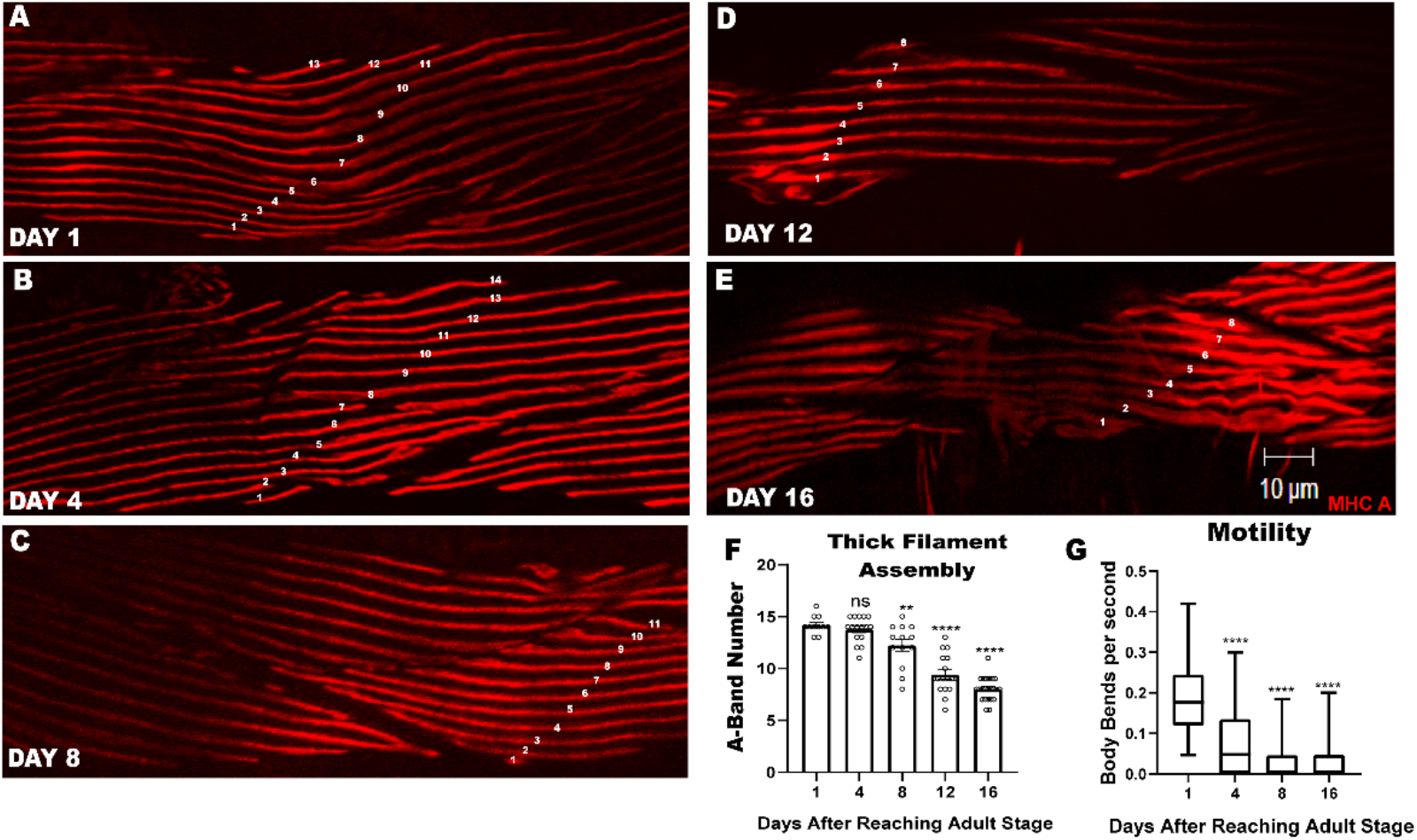
The number of assembled thick filaments (A-bands) and whole nematode motility decline with age. A-E) are representative images of body wall muscle near the vulva immunostained with anti-MHC A at different ages of adulthood (day 1, 4, 8, 12, 16) with an A-band count depicted as white numbers along the A-bands. F) is the quantification of A-band number at different ages of adulthood. G) is the quantification of agar crawling motility assays at different ages of adulthood measured in body bends per second. Statistic depicted are that day of adulthood (4,8,12, or 16) compared to day 1 of adulthood. There was no statistical differences between days 8 and 16 motility. **p-value < 0.005, ***p-value < 0.0005**** p-value < 0.0001.

### UNC-45 has a role in maintaining assembled thick filaments and nematode motility during adulthood

To assess the importance of UNC-45 within *C. elegans* adult muscle we utilized the canonical *unc-45(e286)* temperature sensitive mutant. This strain is relatively normal when grown at 15°C, but has a mutation (L822F) within the UCS domain of UNC-45 that renders it non-functional/reduced when grown at 25°C. When allowed to develop at 25°C, this strain has lower steady state levels of both thick filament isoforms of myosin heavy chain, MHC A and MHC B, and fewer assembled thick filaments (Landsverk et al., 2007; Miller et al., 1983), similar to aged worms. We have shown, using our own antibody to UNC-45, that this mutant does in fact have significantly reduced levels of UNC-45 when grown at 25°C (Moncrief et al., 2021). For our purposes, we allowed the animals to develop normally at 15°C and transferred them to the restrictive temperature of 25°C only after muscle maturity had been reached (referred to as day 0 of adulthood). We observed a decline in assembled thick filaments (A-band number) beginning at day 4 of adulthood assessed by anti-MHC A staining (Figure 2A), as opposed to day 8 in the wildtype strain (Figure 1F). The animals grown at the restrictive temperature also show a decline in motility at day 1 of adulthood (Figure 2B), as opposed to day 4 in both the wildtype (Figure 1G) and the mutant grown at the permissive temperature (Figure 2B). As shown in Figure S2, the growth of wild type animals at 25° does not significantly affect worm locomotion compared to worms growth at 15° when sampled at numerous time points from day 1 to day 16 of adulthood. Within 24 hours of being transitioned to the restrictive temperature perturbed thick filaments can be seen via immunostaining of MHC A in *unc-45(e286)(*Figure 2C-J). We also observed that disorganization of A-bands detected with anti-MHC B staining, also is first apparent at day 4 of adulthood (Figure S2). We would describe this phenotype as an early onset of sarcopenia in *unc-45(e286)* mutant animals. This provides evidence that UNC-45 is essential to muscles during adulthood.

**Figure 2.**
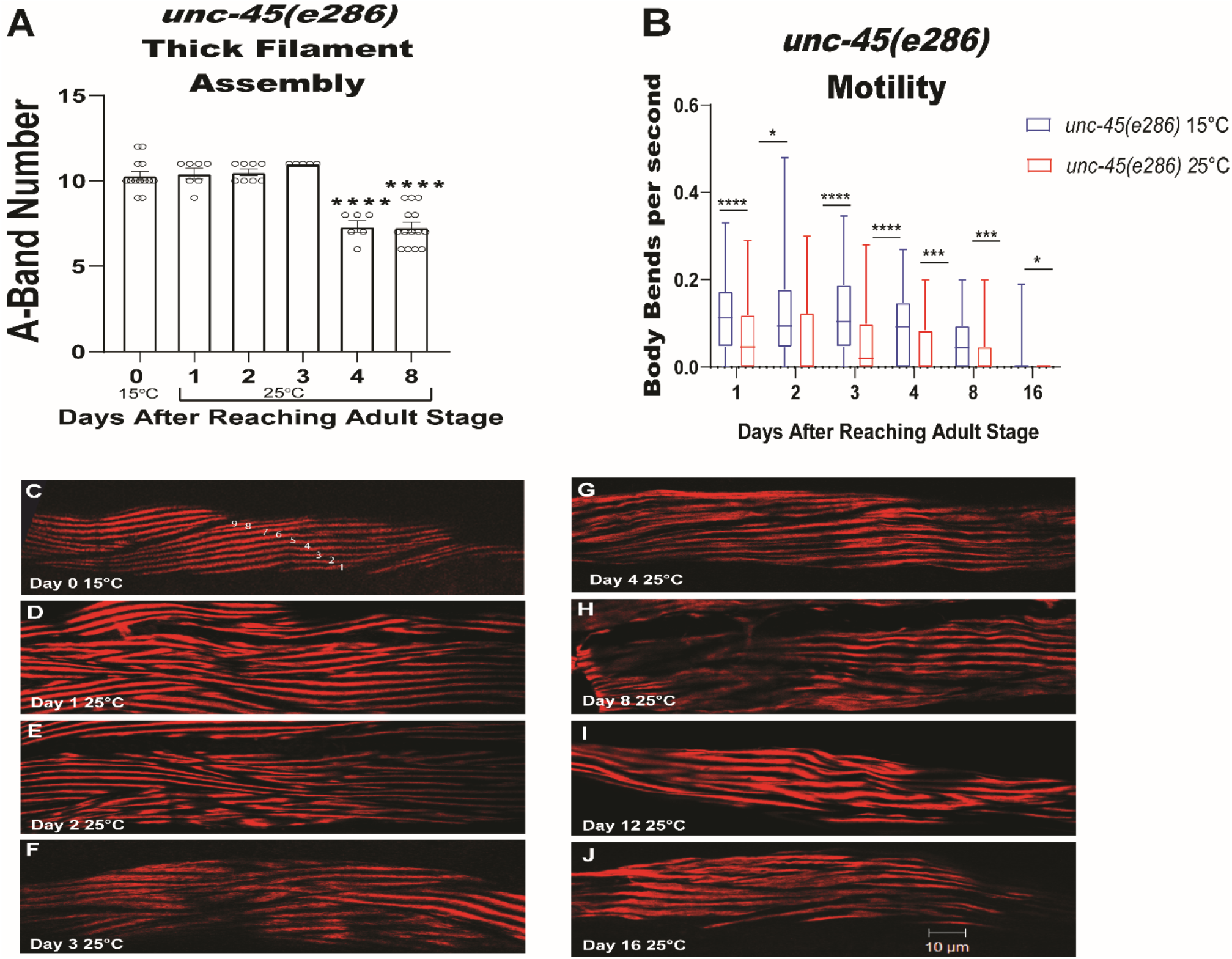
UNC-45 has a role in maintaining assembled thick filaments (A-bands) and nematode motility during adulthood. The canonical *unc-45* temperature sensitive mutant, *e286*, was allowed to develop normally at the permissive temperature of 15°C and shifted to the restrictive temperature of 25°C on day 0 of adulthood. A) is the quantification of A-band number at different ages of adulthood. B) is the quantification of crawling motility assays at different adult ages at the permissive (15°C) and restrictive (25°C) temperatures. C) is a representative image of body wall muscle immunostained with anti-MHC A at day 0 of adulthood after the animal was allowed to develop at 15°C. D-J) are representative images of body wall muscle from animals grown at 25°C immunostained with anti-MHC A at different adult ages (day 1, 4, 8, 12, 16) with an A-band count depicted as white numbers for cells with parallel thick filaments (D-G). **p-value < 0.005, ***p-value < 0.0005**** p-value < 0.0001.

### As adults age, there is a sequential decline of HSP-90, UNC-45 and myosin

Using quantitative western blotting we have found that at day 3 of adulthood, there is a drop in the level of HSP-90 protein (Figure 3A). At day 4 of adulthood there is a drop in the level of UNC-45 protein (Figure 3B). By day 8 of adulthood, there is a drop in the level of MHC B protein, the major client of UNC-45 (Figure 3C). The level of MHC A protein shows a more gradual decline beginning at day 1 (Figure 3D). Although the decline in the level of HSP-90 protein at day 3, may be causally related to the decline in *hsp-90* mRNA at day 2 (Figure 3E), the declines in UNC-45 and MHC-B proteins may be more related to the degradation of UNC-45 and MHC B proteins, rather than the declines in their mRNAs: There is a significant decline in *unc-45* mRNA by day 2 (Figure 3 F), and a significant decline in *unc-54* (encodes MHC B) mRNA by day 1 (Figure 3G), and the levels of these mRNAs remain low throughout the remainder of adulthood. It is noteworthy that the loss of UNC-45 protein at day 4 (Figure 3B) correlates with the onset of reduced whole worm locomotion at day 4 (Figure 1G).

**Figure 3.**
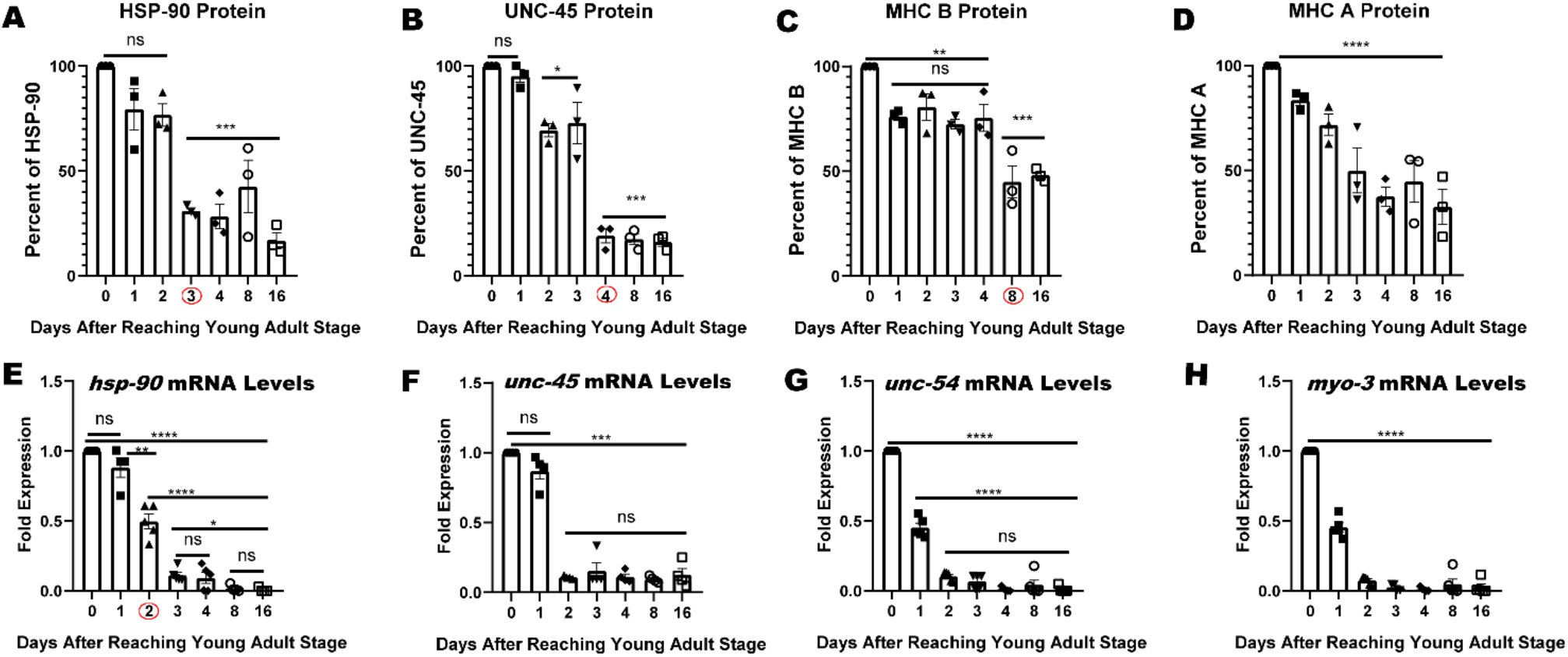
The Sequential Decline of HSP-90, UNC-45, and Myosin with Age. A-D) Graphical quantification of steady state protein levels of HSP-90, UNC-45, MHC B, and MHC A (myosin isoforms). Data are shown as a percentage of protein relative to Histone H3 protein. E-H) Steady state mRNA fold expression of *unc-45, hsp-90, unc-54* (MHC B), and *myo-3* (MHC A) during aging relative to *gpdh-2* (GAPDH). Days of significant protein or mRNA decline are circled with red circles. * p-value < 0.05 **p-value < 0.005, ***p-value < 0.0005**** p-value < 0.0001

### UNC-45 phosphorylation increases with aging

Using gradient SDS-PAGE gels, we noticed that during adult aging, as the protein band corresponding to UNC-45 (107 kDa) declines, a higher molecular weight band appears and increases (Figure. 4). We suspect that this higher molecular weight UNC-45 protein band is the result of post-translational modification. In an attempt to identify this post translational modification, we ran protein lysates from different aged wildtype worms on a SuperSep™ Phos-tag™ electrophoresis gel, which separates proteins based on both molecular weight and phosphorylation status (more phosphate groups cause the protein to run slower through a gel containing the Phos-tag organic molecule which binds to phosphates, including phosphorylated proteins). Using this gel followed by western blotting and reaction with anti-UNC-45 antibodies shows that with age there is an increase in UNC-45 protein bands that run more slowly in the gel beginning at day 3 of adulthood, consistent with a increase in the phosphorylation of UNC-45 with aging (Figure 5A). Further evidence that these slower running UNC-45 bands are phosphorylated was provided by showing that they are eliminated upon treatment of the protein lysates with lambda phosphatase (Figure 5B).

**Figure 4.**
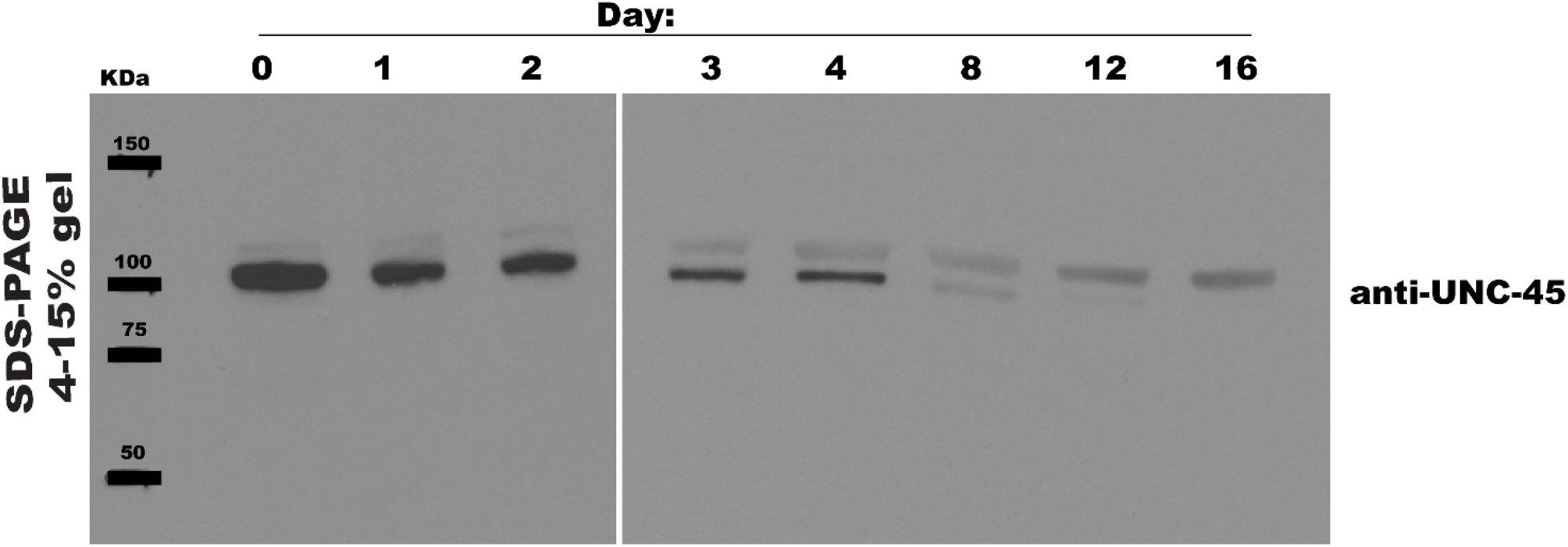
A higher molecular weight UNC-45 band accumulates as the expected sized UNC-45 band declines with aging. Protein lysate from day 0, 1, 2, 3, 4, 8, 12, and 16 old worms were run on a 4-20% gradient gel, transferred to a nitrocellulose blot, and reacted with the UNC-45 antibody to reveal a higher molecular weight UNC-45 band.

**Figure 5.**
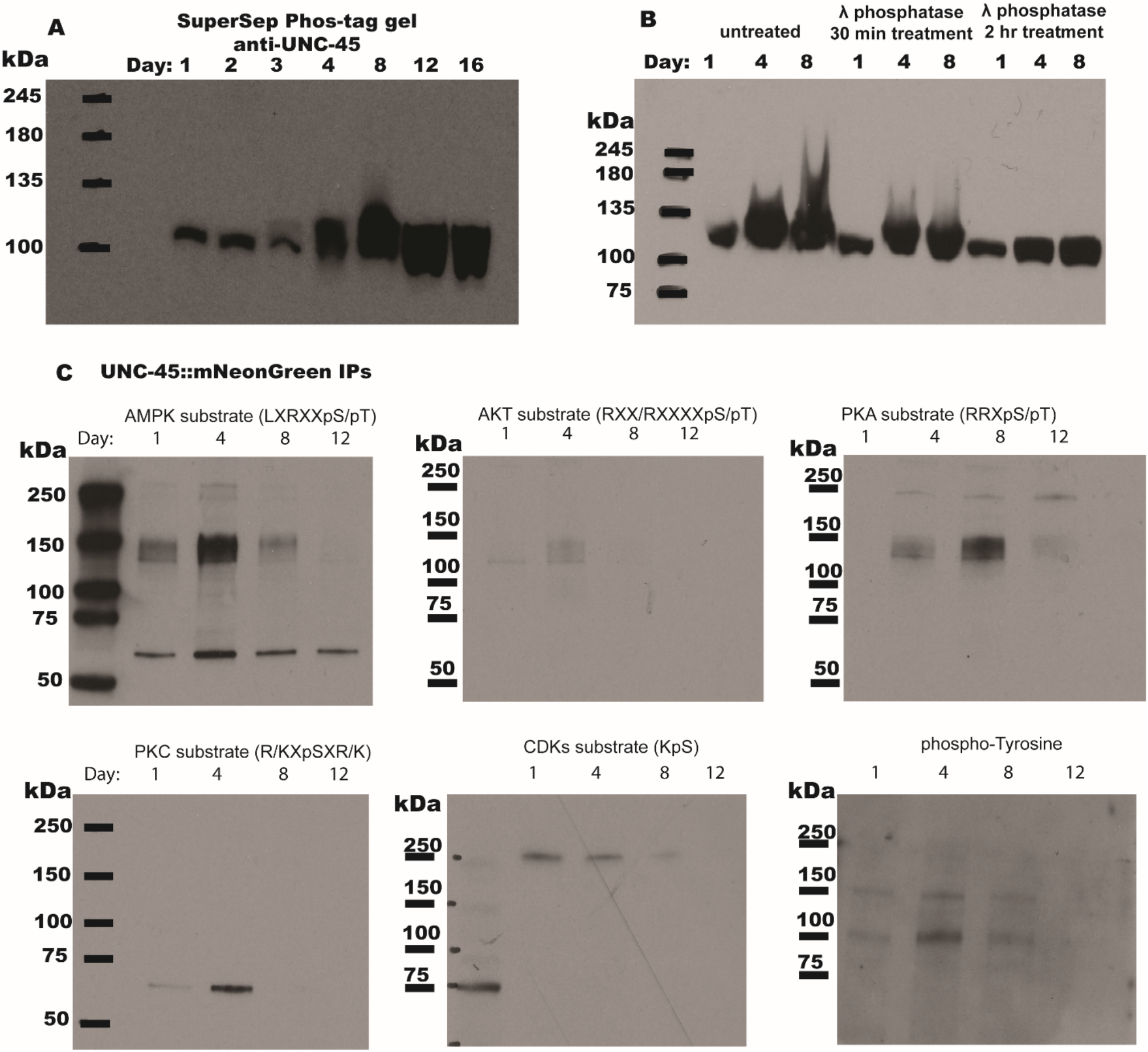
UNC-45 phosphorylation increases with age. A) Wildtype samples at different ages run on SuperSep Phos-tag gels and blotted with anti-UNC-45. The first blot (left) shows the change in phosphorylation of UNC-45 from day 1 to 16 in wildtype animals (7.5% gel). B) shows the putative phosphorylation pattern disappearing after treatment with λ phosphatase (12% gel). C) UNC-45 was immunoprecipitated from CRISPR generated *unc-45::mNeonGreen* worms using nanobodies to mNeonGreen pre-conjugated to magnetic beads and the elutions were run on SDS-PAGE and blotted with different anti-phospho antibodies specific for different phosphorylation motifs.

We also immunoprecipitated endogenous UNC-45::mNeonGreen from a CRISPR generated strain using magnetic beads coupled to anti-mNeonGreen nanobodies, and reacted the eluted bound fraction with different anti-phospho-antibodies. We found increased phospho-serine/threonine and phospho-tyrosine at day 4 of adulthood, which is when we observe the dramatic decline of UNC-45. Based on the phospho-antibodies and the motifs they are reported to react with, UNC-45 is potentially being phosphorylated by multiple kinases, including PKC, AMPK, PKA, and AKT, which all recognize similar phosphorylation motifs. Additional phosphorylated protein(s) appear to be pulled down with the UNC-45::mNeonGreen immunoprecipitation and react with the phospho-antibodies (Figure 5C). The band running around 250kDa is likely one or more Myosin isoforms, and the band running around 60kDa could be HSP-70, which is known to complex with UNC-45(Barral et al., 2002; Liu et al., 2008; Srikakulam and Winkelmann, 2004). To identify the site or sites of phosphorylation on UNC-45, we immunoprecipitated UNC-45::mNeonGreen from a large quantity of worms of mixed developmental stages, including adults of different ages. This UNC-45::mNeonGreen was subjected to mass spectrometry analysis. After cleavage with trypsin and GluC, peptides covering about 94% of the UNC-45 protein sequence were analyzed and the only site of phosphorylation identified was serine 111. As shown in Figure 6, this residue lies close to the C-terminus of the TPR domain, and could conveivably interfere with binding to the HSP-90 C-terminus. Intriguingly, serine 111, though in a highly conserved region, is not a conserved residue. In fact, in Drosophila it is an aspartic acid and in humans and zebrafish it is a glutamic acid – both negatively charged residues. Perhaps the negative charge here is important and the residue evolved to have an inherent negative charge.

**Figure 6.**
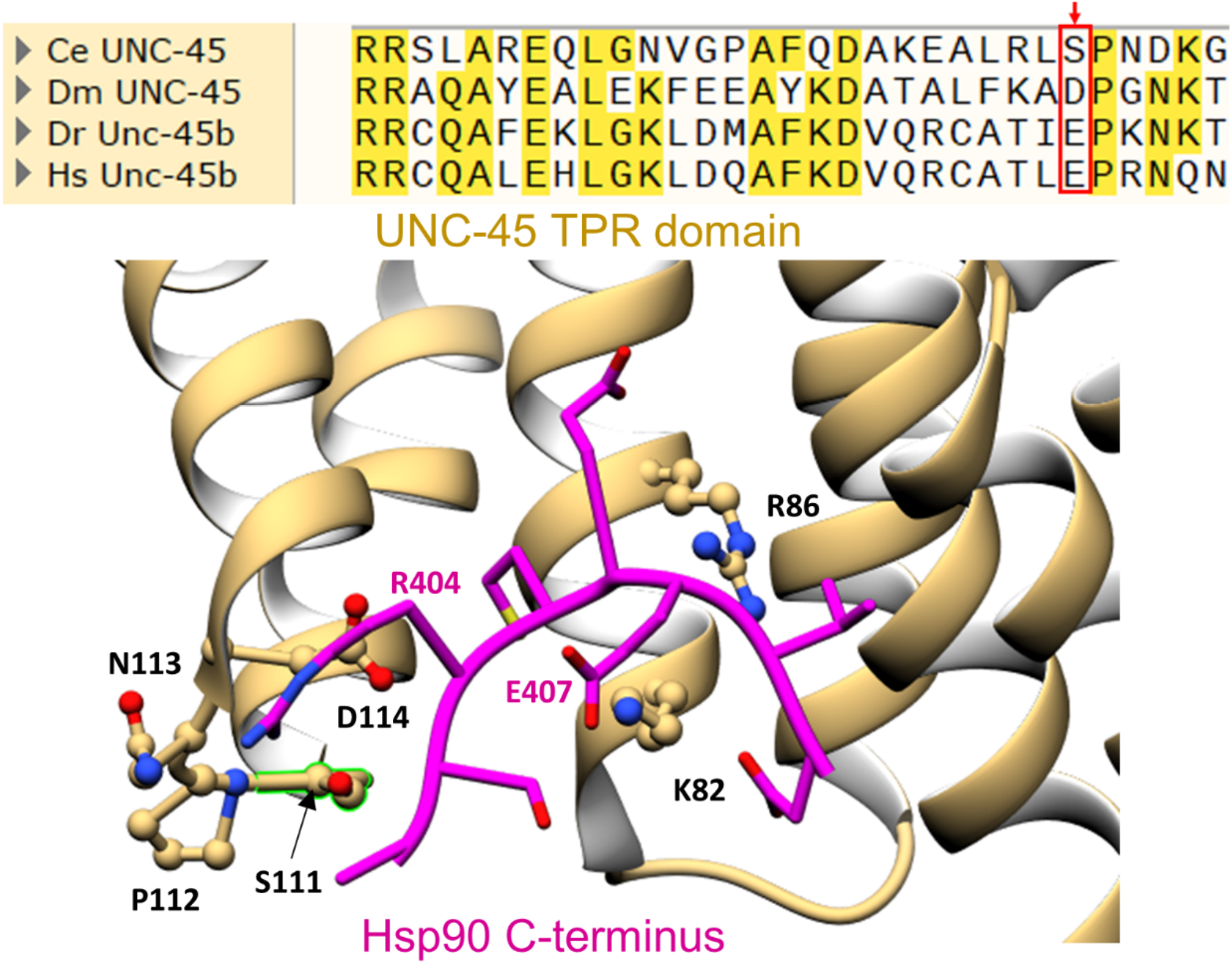
UNC-45 is phosphorylated at Serine 111. Sequence alignment of *C. elegans, Drosophila*, zebrafish, and human UNC-45/Unc-45b at the end of the TPR domain. The site of *C. elegans* phosphorylation, S111, is indicated with a red arrow and the sequence alignment is boxed in red. Below the alignment is the crystal structure of the UNC-45 – HSP-90 interaction site in *C. elegans*, with the UNC-45 TPR domain in yellow, and the HSP-90 C-terminus in purple (modified from the crystal structure of C. elegans UNC-45 that includes an HSP-90 C-terminal pepetide, 4i2z.pdb (Gazda et al., 2013)).

### A delayed onset of sarcopenia is associated with increased UNC-45 and HSP-90

It is well-established that genetic disruption of the insulin-like signaling pathway results in increased longevity of *C. elegans* (Kenyon et al., 1993; Klass, 1983). The *age-1(hx546)* strain has a mutation in the phosphatidylinositol 3-kinase catalytic subunit (PI3KCA) of the insulin like signaling pathway that results in animals living ∼7-9 fold longer (Ayyadevara et al., 2008). We have found that these animals experience a delayed onset of sarcopenia, with the number of assembled thick filaments only marginally declining by day 16 of adulthood (Figure 7F). Intriguingly, although it has been reported that these animals move more in liquid media early in life (Duhon and Johnson, 1995), we found a slight decline in spontaneous movement on agar plates at days 1 and 4 (Figure 7G). This does not necessarily correlate to muscle health as it could be due to uncharacterized neuronal or metabolic changes caused by this mutation. When stimulated by poking with a toothpick, the animals display normal, healthy sinusoidal movement on the agar surface until at least day 8 of adulthood. These mutant animals not only retain their HSP-90, UNC-45 and MHC B protein levels longer than wildtype, but the steady state levels of these proteins continue to increase past day 0 of adulthood (Figure 8A-C). This is not surprising since we observe that the steady state transcripts of *hsp-90* and *unc-54* continue to increase past day 0 of adulthood, spiking around days 2 and 3, as if the animals are still developing. In contrast, we don’t see much change in *unc-45* transcript levels between wildtype and the long-lived animals. We once again found that the decline in UNC-45 protein occurs independently of its transcript and after the decline in HSP-90 protein.

**Figure 7.**
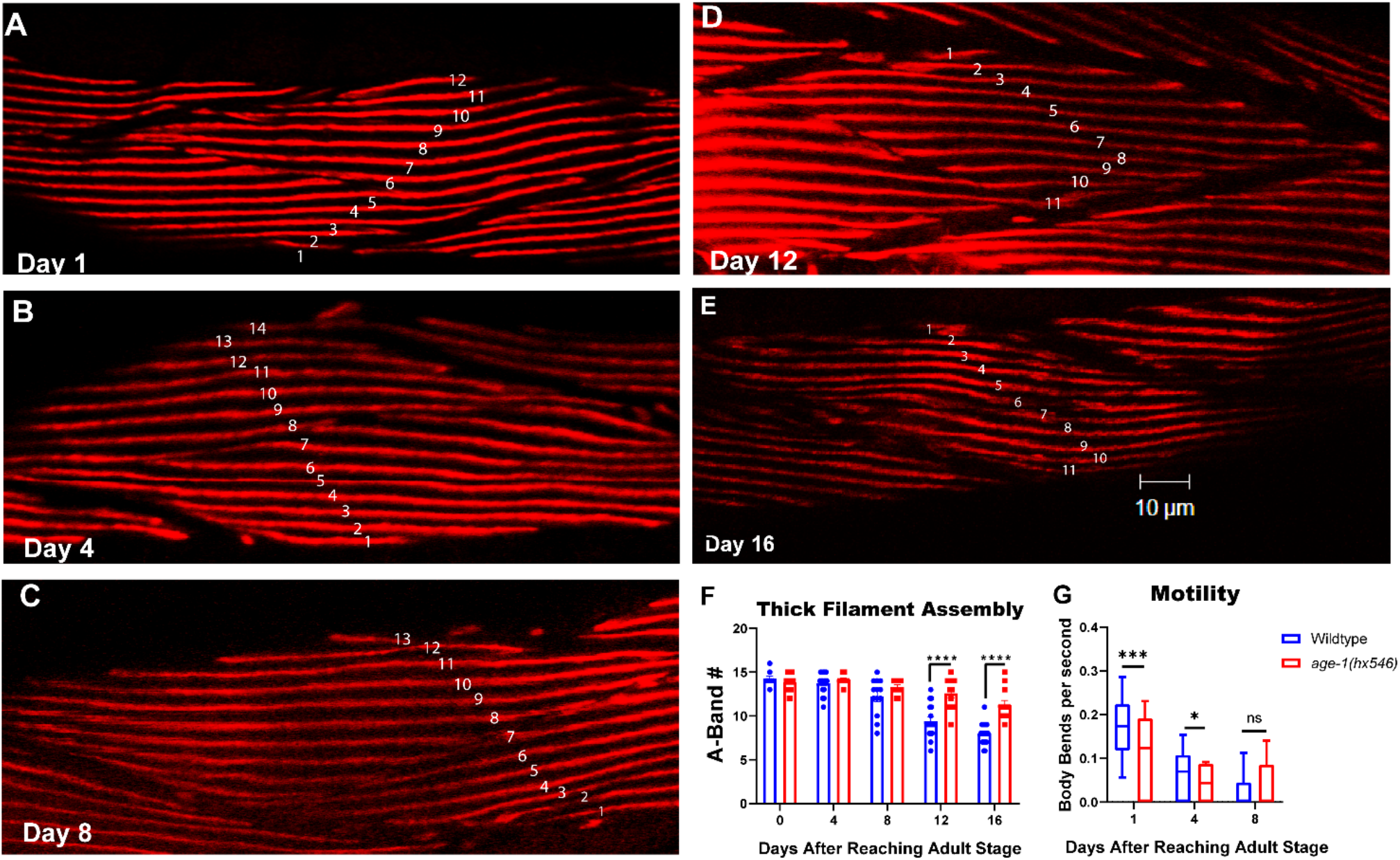
The *age-1(hx546)* longevity mutant has a delayed loss of assembled thick filaments but not motility. A-E) are representative images of body wall muscle near the vulva immunostained with anti-MHC A at different ages of adulthood (day 1, 4, 8, 12, 16) with an A-band count depicted as white numbers along the A-bands. F) is the quantification of A-band number at different ages of adulthood compared to N2 wildtype. G) is the quantification of agar crawling motility assays at different ages of adulthood measured in body bends per second compared to N2 wildtype. **p-value < 0.005, ***p-value < 0.0005 **** p-value < 0.0001.

**Figure 8.**
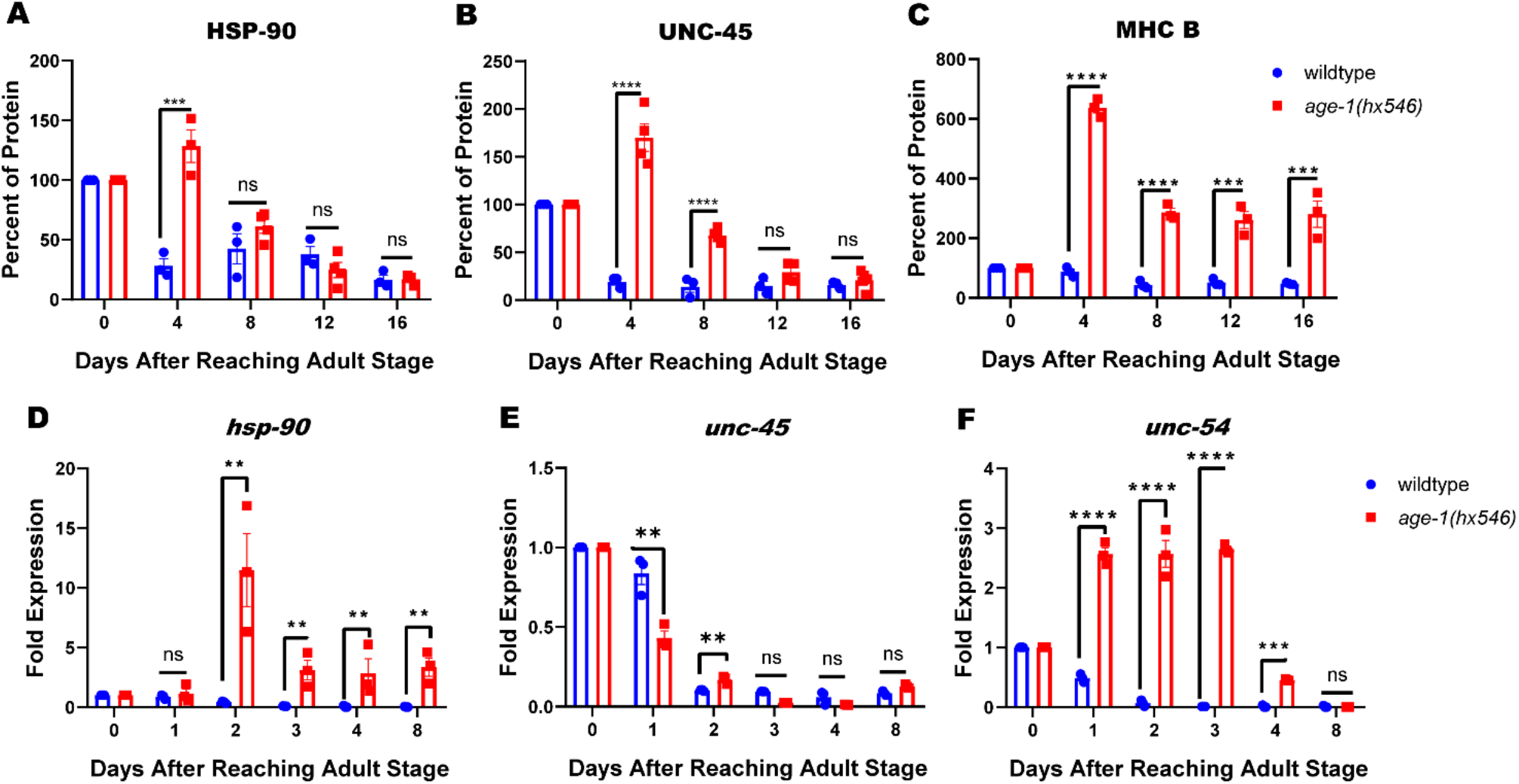
The *age-1(hx546)* longevity mutant has increased levels of UNC-45, HSP-90, and myosin MHC-B. A-C) Graphical quantification of steady state protein levels of HSP-90, UNC-45, and MHC B (major body wall myosin isoform). Data are shown as a percentage of protein relative to histone H3 protein. D-F) Steady state mRNA fold expression of *unc-45, hsp-90* and *unc-54* (MHC B) during aging relative to *gpdh-2* (GAPDH). * p-value < 0.05 **p-value < 0.005, ***p-value < 0.0005**** p-value < 0.0001

### The interaction of HSP-90 with UNC-45 may stabilize UNC-45

The TPR domain of UNC-45 binds to HSP-90 (Barral et al. 2002). Intriguingly, we have found that the *hsp-90(p673)* temperature sensitive mutant has significantly less UNC-45 protein when grown at the restrictive temperature of 25°C (Figure 9B and D). This loss of protein appears to be independent of steady state mRNA since *unc-45* mRNA increases at the restrictive temperature of 25°C (Figure 8E). This also provides evidence that *unc-45* transcription may be sensitive to heat stress, and potentially other types of stress. The *hsp-90(p673)* mutation (E292K) is described as a weak gain of function mutation and speculated to potentially disrupt interactions with some clients and/or co-chaperones(Birnby et al., 2000). The mutation is located on the surface of the middle domain (MD) of the protein (Figure 9A), while the ATPase region is in the N-terminal domain (NTD)(Obermann et al., 1998) and UNC-45 binds to the conserved MEEVD sequence towards the end of the C-terminal domain (CTD)(Barral et al., 2002). Because this mutant HSP-90 results in such a dramatic loss of UNC-45 protein, we speculate that this mutation is somehow affecting HSP-90’s ability to interact with UNC-45. However, since the known binding site of HSP-90 in its C-terminus (CTD) to UNC-45 is so distant from the mutation site in its MD, perhaps the mutation causes some conformational change that is transmitted from the MD to the CTD. The *unc-45(e286); hsp-90(p673)* double mutant displays reduced motility to that of *unc-45(e286)* alone at the permissive temperature of 15°C (Figure 9F). This would suggest that the additional mutation is enough to exacerbate the Unc phenotype even at the permissive temperature. Above 15°C, all three strains exhibit little to no crawling motility.

**Figure 9.**
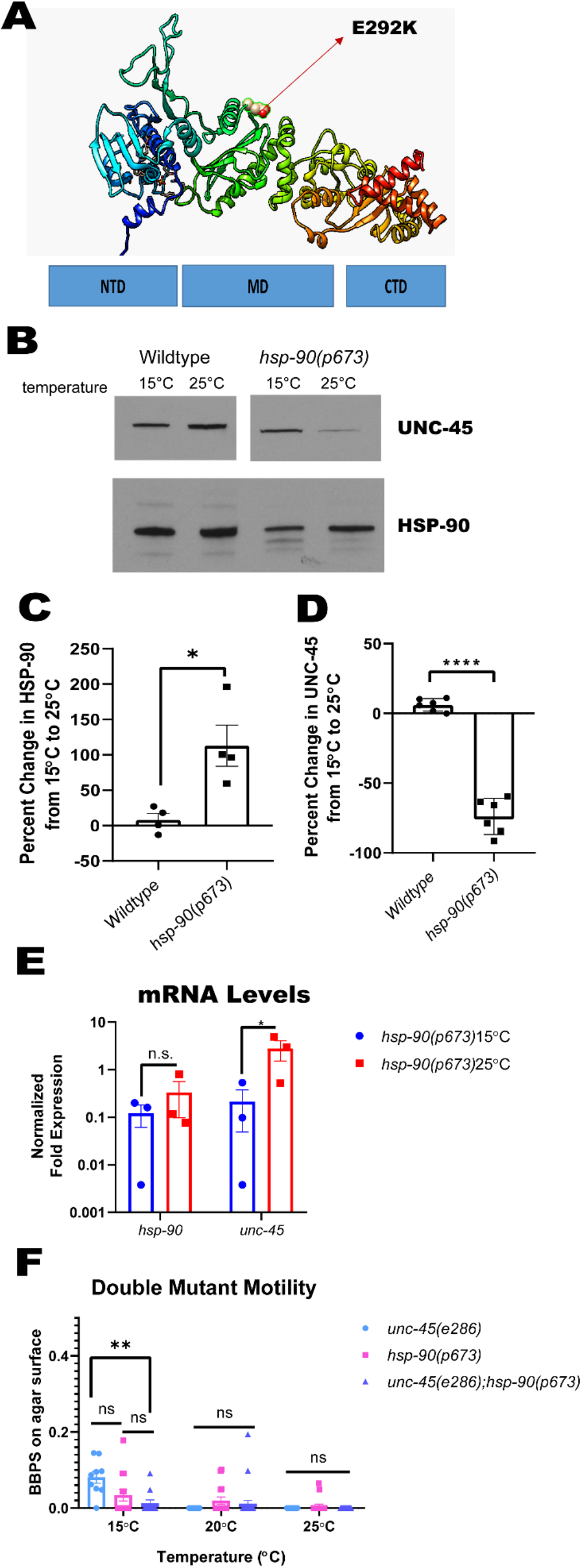
*hsp-90* loss of function temperature sensitive mutant has decreased UNC-45 protein, but not transcript. A) A homology model of the nematode HSP-90 protein showing the location of the E292K mutation in *hsp-90*(*p673*). B) Western blot of UNC-45 and HSP-90 steady state protein levels. Histone H3 was used as the protein loading control. C and D) Quantification of UNC-45 and HSP-90 protein percent reduction at 25°C relative to 15°C from wild type and *hsp-90(p673)*. E) The relative fold expression of *unc-45* and *hsp-90* mRNA levels of the *hsp-90(p673)* strain grown at 15°C and 25°C. *ges-1* (gut esterase) was used to normalize expression. F) The quantification of agar crawling motility of the *unc-45(e286), hsp-90(p673)*, and *unc-45(e286);hsp-90(p763)* measured in body bends per second mutant strains. * -value < 0.05 **p-value < 0.005, ***p-value < 0.0005**** p-value < 0.0001.

## Discussion

With conserved pathways, a short lifespan, large sample sizes, and a plethora of easy genetic techniques available, *C. elegans* have become a vital model organism in the pursuit of studying the molecular mechanisms responsible for many aging associated pathologies, including sarcopenia. With age, we begin to lose our ability to generate new muscle cells due to the dwindling population of satellite stem cells. This makes the maintenance of already existing muscle fibers key to maintaining muscle health with age. With this in mind, the nematode becomes an even more intriguing model to use for sarcopenia studies since they do not have satellite cells to replenish their muscle cells. In this study we have used *C. elegans* to identify changes in the myosin head chaperone UNC-45 during aging that may prove crucial to the decline in muscle maintenance with age.

Previously, UNC-45 had only been shown to be crucial during muscle development, but not during adulthood (Epstein and Thomson, 1974). However, since then more evidence has emerged that suggests that myosin may require its chaperone UNC-45 to maintain its integrity in the face of damage that accumulates with age. We now know that sarcomere proteins assembled into the thick filament have a low protein turnover rate(Solomon and Goldberg, 1996) and it has been shown that the reduction in myofibrillar synthesis rate in old (61-74yr) vs. young (22-31yr) human muscle is not caused by reduced myosin mRNA(Welle et al., 1996). This suggests that the myosin assembled into thick filaments requires a mechanism to retain its structure after denaturing events (chemical, thermal, and physical stress) that occur within the muscle. Barral et al. have shown that UNC-45 prevents thermal aggregation of myosin heads in vitro (Barral et al., 2002). Etard et al. have shown that Unc-45b, the skeletal muscle specific isoform, localizes to the Z-discs normally and moves to thick filaments during stress (cold and heat shock, chemical, and physical) in zebrafish skeletal muscle (Etard et al., 2008). We find that perturbation of UNC-45 during adulthood is enough to cause an early onset of sarcopenia-like pathology. Thus, our results demonstrate that UNC-45 is crucial to maintaining muscle health specifically during adulthood.

We then sought to characterize the changes in total steady state protein and transcript levels of UNC-45, its co-chaperone HSP-90, and its clients within body wall muscle, MHC A and MHC B (Figure 3). We found that *hsp-90* transcript declines at day 2 of adulthood, directly before the decline in HSP-90 protein. The loss of HSP-90 at day 3 directly precedes a loss of UNC-45 protein at day 4 of adulthood. Then, there is a major decline in MHC B, the main client of UNC-45 and more abundant body wall muscle myosin isoform, at day 8 of adulthood. We theorize that the decline in UNC-45 protein is due primarily to protein degradation. The decline in UNC-45 that we observe during adulthood may be due to increased ubiquitination followed by proteasomal degradation. It has been reported that *C. elegans* UNC-45 is polyubiquitinated by CHN-1 and CDC-48 resulting in degradation by the 26S proteasome (Hoppe et al., 2004). One study found that aged sarcopenic rat skeletal muscle contains higher levels of CHIP and p97, which are the mammalian homologs of CHN-1 and CDC-48(Altun et al., 2010). The authors found no significant change in *unc-45b* mRNA between young adult and aged muscle samples but found that Unc-45B protein was decreased in the aged muscle compared to young adult muscle(Altun et al., 2010). We observe what appears to be an increase in post-translational modification of UNC-45 with age (Figure 4), and have identified that one of these modifications is likely to be phosphorylation (Figure 5). Phosphorylation may be related to the already known ubiquitylation of UNC-45 and its degradation by the proteasome (Janiesch et al., 2007). Through mass spectrometry, we have identified serine 111 as a site of phosphorylation. Adding a bulky negative charge to this region of UNC-45’s TPR domain could interfere with the interaction between UNC-45 and HSP-90. Though in a conserved region, serine 111 is not a conserved residue. In humans this residue is actually a glutamic acid, which would act like a phosphomimetic. This raises the question of why this residue evolved from being phosphorylated to being consititutively negatively charged. Perhaps this site is crucial to the regulation of UNC-45 protein levels within cells. Whether it interferes with HSP-90 binding or leads to degradation of UNC-45 remains unexplored. A further caveat is that S111 was found to be phosphorylated in UNC-45 purified from a population of mixed developmental stages and adults of various ages. Thus, we do not yet know if S111 is a site that is phosphorylated during muscle aging.

The *age-1(hx546)* longevity mutant shows a delay in the loss of assembled thick filaments but not spontaneous crawling motility (Figure 7). A major characteristic of aging muscle and sarcopenia pathology is the loss of muscle mass, which does not begin to occur in these animals until day 16 of adulthood (day 8 in wildtype). The amount of UNC-45, HSP-90, and MHC B protein continues to increase in these animals well past day 0 of adulthood. *hsp-90* and *unc-54* transcripts remain above wildtype levels while *unc-45* transcript has the same trend as in the wildtype animals. Since one theory behind the improved stress resistance of this strain is an increase in heat shock proteins and chaperones(Shmookler Reis et al., 2012; Walker et al., 2001), it is not surprising that the *hsp-90* mRNA is upregulated. However, we do still observe a decline in HSP-90 protein back to wildtype levels by day 8 of adulthood, which precedes a decline in UNC-45 protein back to wildtype levels by day 12 of adulthood. This delay in the loss of UNC-45 protein may be one of the mechanisms that allow this long lived mutant to have an extended health span as well as lifespan.

Having increased UNC-45 to maintain myosin heads during aging would be beneficial to refold the myosin head after stress if there is also equivalent HSP-90 available to re-bind the TPR domain so as not to cause aberrant inhibition of the myosin power stroke. We find that in a loss of function *hsp-90* mutant there is significantly reduced UNC-45 protein, but not transcript. This suggests that interaction with HSP-90 is important for the protein stability of UNC-45 and that in the absence of HSP-90, UNC-45 is left more susceptible to degradation. This leads us to theorize that during aging there is a loss of HSP-90, which leaves regions of UNC-45 more open to post translational modifications, like phosphorylation, and leads to increased degradation of UNC-45. At least for C. elegans, the crystal structure of UNC-45 (Gazda et al., 2013) shows that it forms linear multimers in which the length of the repeating unit is similar to the repeating unit in the staggered display of pairs of myosin heads on the surface of thick filaments. In addition, antibodies to UNC-45 co-localize with MHC B myosin on the major portions of the thick filament in already assembled sarcomeres of adult muscle (Ao and Pilgrim, 2000; Gazda et al., 2013). Thus, in adult muscle, UNC-45 is postioned on the thick filament to re-fold any myosin heads that are damaged by thermal or chemical stress. Therefore, our model is that during adult aging, the loss of HSP-90, followed by the loss of the myosin head chaperone UNC-45, leads to fewer myosin heads being re-folded, and these myosin molecules and even thick filaments are removed from the sarcomere, forming aggregates and/or being degraded. Because of the low level of myosin transcripts and translation, there is not enough new myosin protein available to replenish the myosin within existing thick filaments, or to form new thick filaments. This results in an overall decline in thick filament number, sarcomere size and number, and muscle function.

As we age, we experience a loss of our satellite cells (muscle stem cells) and the maintenance of our existing muscle cells becomes crucial to maintaining our mobility. UNC-45 is the conserved myosin head chaperone responsible for the initial folding and assemblage of myosin heads into the thick filament, and *most likely* refolding the myosin head after stress causes it to unfold. Humans have two UNC-45 homologs, A and B. UNC45-A is expressed ubiquitously and UNC-45B is expressed primarily in striated muscles – skeletal and cardiac. The results presented here suggest that increased expression and/or increased activity of UNC45B might be a strategy for prevention and treatment of sarcopenia, and even age-associated heart failure.

## Materials and Methods

### *C. elegans* Strains

Standard growth conditions for *C. elegans* were used (Brenner, 1974). Wildtype Bristol N2, *age-1(hx546), GB319, and GB350* were grown at 20°C. As described in the above results, temperature sensitive mutants *unc-45(e286)* and *hsp-90(p673)* were grown at 15°C, 20°C, or 25°C. Adult worms were separated from their progeny daily by allowing the adults to sink in M9 buffer in glass tubes, removing the supernatant containing the L1s, and washing several times before returning to NGM plates. The double mutant *unc-45(e286); hsp-90(p673)* was generated by genetic crossing. GB319 is the 2X outcrossed derivative of PHX789 (*unc-45(syb789))* which is a CRISPR-generated strain that expresses UNC-45-mNeonGreen and was described previously (Moncrief et al., 2021). The strain, PHX501 (*hsp-90(syb501))*, is a CRISPR-generated strain which expresses HSP-90 with a C-terminal mKate2 tag. PHX501 was created by SunyBiotech (http://www.sunybiotech.com). PHX501 was outcrossed 2X to wild type to generate strain GB350.

### Immunostaining in adult body-wall muscle

Adult nematodes were fixed and immunostained according to the method described by Nonet et al. and described in further detail by Wilson et al. (Nonet et al., 1993; Wilson et al., 2012). The following primary antibodies were used: anti–MHC A at 1:200 (mouse monoclonal 5-6; Miller et al., 1983), anti-MHC B at 1:200 (mouse monoclonal 5-8 (Miller et al., 1983)), and anti–UNC-95 at 1:100 (rabbit polyclonal Benian-13 (Qadota et al., 2007)). Secondary antibodies, used at 1:200 dilution, included anti-rabbit Alexa 488 (Invitrogen, Thermo Fisher Scientific) and anti-mouse Alexa 594 (Invitrogen). Images were captured at room temperature with a Zeiss confocal system (LSM510) equipped with an Axiovert 100M microscope and an Apochromat ×63/1.4 numerical aperture oil immersion objective in ×2.5 zoom mode. The color balances of the images were adjusted by using Photoshop (Adobe, San Jose, CA).

### Motility crawling assays

Adult worms were collected at different ages using M9 buffer containing 0.2g/L gelatin. They were transferred to a 1.5mL microcentrifuge tube, allowed to settle to the bottom, and washed 3X with M9 buffer containing 0.2g/L gelatin. 5µL of worm suspension was added to the center of a 6cm unseeded NGM plate. Worms were allowed to adapt for 5 minutes before a video recording of their crawling was made using a dissecting stereoscopic microscope fitted with a CMOS camera (Thorlabs). Several 10 second videos were recorded for each sample and analyzed by Image J FIJI WrmTracker software to obtain body bends per second (BBPS). Statistical significance was determined using a student’s T-test.

### Quantitative Real-Time PCR

Worms from two 10cm NGM plates were collected using M9 buffer and frozen as a dry packed worm pellet. Lysis buffer from the Qiagen RNeasy Plus Mini Kit (cat. 74134) was added, the sample was freeze-thawed in liquid nitrogen five times to crack the worm cuticle, and then vortexed with MagnaLyser beads (Roche) for 1 minute. The samples were centrifuged, and the supernatant was used to extract RNA with the Qiagen RNeasy Plus Mini Kit. The NCBI primer designing tool (https://www.ncbi.nlm.nih.gov/tools/primer-blast/) was used to create primer pairs specific for each gene of interest that also had at least one intron separating the primer pair (see Table S1). 1µg of cDNA was synthesized from the RNA using BioRad iScript Reverse Transcription Supermix (cat. 1708840). The cDNA was then diluted 1/10 in nuclease free water and 2.5µL was used per reaction with 10µL of qRT-PCR master mix (1.25µL of each primer, 1.25µL of nuclease free water, and 6.25 µL of SybrGreen supermix (Biorad cat. 1708880)). A BioRad CFX Real-Time PCR thermal cycler was used. Fold change was determined using the 2^ddCt method (Schmittgen and Livak, 2008). *ges-1* and *gapdh-2* were used for normalization. Statistical siginifcance was determined using a student’s T-test.

### Production and purification of antibodies

The generation of rabbit polyclonal antibodies to the C-terminal 120 residues of UNC-45 was described previously (Moncrief et al., 2021). A 149 residue region (aa 553-702) at the C-terminus of HSP-90 (“HSP-90 antigen”)(Figure S4A) was expressed in E. coli as a GST fusion protein and sent to Noble Life Sciences for antibody production in rats. After approx. 3 months, we received the antisera, and affinity purified anti-HSP-90 antibodies by use of an MBP-HSP-90 antigen coupled to Affigel matrix (BioRad), using a procedure previously published (Benian et al., 1993). Both antibodies work well in western blots, detecting a polypeptide of expected size for UNC-45 (107 kDa) and HSP-90 (80 kDa), and no detectable extraneous bands at a 1:5,000 or 1:2,500 dilution, respectively. In addition, we demonstrated that anti-HSP-90 antibodies detect a protein of expected size of approximately 100 kDa from a lysate prepared from animals that express HSP-90-mKate2 (Figure S3B).

### Western blots and quantitation of protein levels

We used the procedure of Hannak et al. (Hannak et al., 2002) to prepare total protein lysates from wild-type, *hsp-90(p673)*, and *age-1(hx546)* strains. When comparing wild-type and mutant strains, we loaded approximately equal amounts of protein extract estimated by finding volumes of extracts that would give equal intensity of banding after Coomassie staining. We used quantities of extracts and dilutions of antibodies that would place us into the linear range of detection by ECL and exposure to film. The following antibodies and dilutions were used: rabbit anti–UNC-45(Moncrief et al., 2021) at 1:5,000; rat anti–HSP-90 at 1:2,500; mouse monoclonal 5-8 (Miller et al., 1983) for MHC B at 1:40,000; mouse monoclonal 5-6 ascites (Miller et al., 1983) for MHC A at 1:5,000; Phospho-(Ser/Thr) Kinase Substrate Antibody Sampler Kit (cell signaling) at 1:1,000; Phospho-Tyrosine (4G10, cell signaling) mouse mAb at 1:1,1000. The quantitation of steady-state levels of protein was performed as described in Miller et al. (2009)(Miller et al., 2009). The relative amount of each of these muscle proteins in each lane was normalized to the amount of Histone H3 detected using anti-Histone H3 (abcam, cat. ab1791) at 1:40,000 dilution. The amount of Histone H3 at different ages was compared to Ponceau S staining to ensure no significant changes throughout aging (Figure S5). BioRad precast Mini-PROTEAN TGX Stain-Free Gels were used (4-20%, cat. 4568093, 12%, cat. 4568041). The BioRad Precision Plus Protein™ Kaleidoscope™ Prestained Protein Standards (cat. 1610375) were used on all standard SDS-PAGE gels.

### Immunoprecipitation

Worms were collected from 2 to 4 10cm NGM plates with M9 and frozen. 500µL – 1mL of IP buffer (25mM Tris pH 7.5, 150mM NaCl, 1mM EDTA, 0.5% NP40, 5% glycerol, 1X Halt protease & phosphatase inhibitor cocktail (ThermoFisher)) was added to each sample. They were then freeze-thawed in liquid nitrogen 3-5X to crack the cuticle, added to MagnaLyser beads, and vortexed for 1 minute. The samples were centrifuged, and the lysate supernatant was added to 25µL of mNeonGreen-Trap Magnetic Agarose suspension (cat. no. ntma-10, Chromotek, Inc.) in which nanobodies to mNeonGreen had been coupled to magnetic agarose beads. They were then incubated on a spinning wheel at 4°C for 1 hour, washed 3X with IP buffer, and eluted with 2X Laemmelli buffer with βME.

### Phosphorylation analysis

SuperSep™ phos-tag™ SDS gels (Fuji Film 198-17981, 195-17991) were used per product instructions. Samples were prepared using the IP method described above and cleaned with Amicon centrifugal filters. The WIDE-VIEW™Prestained Protein Size Marker III (Fuji Film, 230-02461) was used with all SuperSep™ phos-tag™ SDS gels. Lambda protein phosphatase (New England BioLabs, P0753S) was used per product instructions.

### Mass Spectrometry Sample Preparation

Samples were immunoprecipitated from 3 generous scoops of worm powder (extensively ground in a mortar and pestle in liquid nitrogen) made from a mixed population of UNC-45::mNeonGreen worms using the above immunoprecipitation method. Initially, the elution was run on a 4-20% SDS-PAGE, and the resulting bands were excised and sent to the mass spectrometry facility at the University of Texas Medical Branch (Galveston, Texas), where they were analyzed by Dr. Aaron Bailey. Because of the purity of the initial samples, we then sent Dr. Bailey the in solution elution to analyze.

### Nanoflow-LC-MS/MS peptide mapping

Immunoprecipitated UNC-45-mNG samples were prepared for LC-MS/MS analysis as previously described (Su et al., 2020). Samples were analyzed by nanoLC-MS/MS (nanoRSLC, ThermoFisher) using an Aurora series (Ion Opticks) reversed phase HPLC column (25 cm length × 75 µm inner diameter) directly injected to an Orbitrap Eclipse using a 120 min gradient (mobile phase A = 0.1% formic acid (Thermo Fisher), mobile phase B = 99.9% acetonitrile with 0.1% formic acid (Thermo Fisher); hold 12% B for 5 min, 2-6% B in 0.1 min, 6-25% in 100 min, 25-50% in 15 min) at a flow rate of 350 nL/min. Eluted peptide ions were analyzed using a data-dependent acquisition (DDA) method with resolution settings of 120,000 and 15,000 (at *m/z* 200) for MS1 and MS2 scans, respectively. DDA-selected peptides were fragmented using stepped high energy collisional dissociation (20%, 30%, 40%). Tandem mass spectra were analyzed according to a label-free proteomic strategy using Proteome Discoverer (version 2.5.0.400, ThermoFisher) with the Byonic (version 4.1.10, Protein Metrics) and Minora nodes using the unreviewed C. elegans proteome (Uniprot, downloaded 6 May 2021) and the Unc45-mNG sequence as a reference FASTA database(Bern et al., 2012; Lin et al., 2018). Mass tolerances of 10 ppm and 20 ppm were used for matching parent and fragment masses, respectively. Mass spectra were searched with a fixed modification of carbamidomethyl (C), and up to 2 common variable modifications of deamidation (N,Q), oxidation (M), phosphorylation (S, T, Y), with Wildcard Search enabled, allowing for a mass tolerance of −40 Da to +400 Da. Peptide spectral matches were filtered for quality: PEP2D < 0.01, Byonic score > 100 (Tian et al., 2021).

### Homology modeling of nematode HSP-90

For HSP-90 protein structure modeling, CLUSTALW version 1.2.2 (https://www.ebi.ac.uk/Tools/msa/clustalw2/), SWISS-MODEL version July 2021 (https://swissmodel.expasy.org/;(Waterhouse et al., 2018)) online tools were used. Human Hsp-90 (5fwm.pdb; (Verba et al., 2016)) was used as template crystal structure and the *C. elegans* HSP-90 sequence as the target. Molecular graphics were generated by using Chimera version 1.15 (https://www.cgl.ucsf.edu/chimera/; (Pettersen et al., 2004))

## Acknowledgments

This work was supported in-part by National Institutes of Health grant R01GM118534 to G.M.B. and A.F.O., and also supported in-part by the Emory Initiative Biological Discovery Through Chemical Innovation (BDCI) to G.M.B. Many of the nematode strains used in this work were provided by the Caenorhabditis Genetics Center, which is funded by the NIH Office of Research Infrastructure Programs (P40 OD010440). The Mass Spectrometry Facility at UTMB is supported in part by Cancer Prevention Research Institute of Texas (CPRIT) grant number RP190682. Monoclonal culture supernatants for 5-6 and 5-8 antibodies were obtained from the University of Iowa Hybridoma Bank, and the 5-6 ascites fluid was kindly provided by Henry F. Epstein (deceased).

## Author Contributions

C.J.M., H.Q. and G.M.B. conceived the study. C.J.M. performed most of the experiments, their analysis and interpretation. A.O.B. performed and interpreted the mass spec analysis. M.K. helped perform some of the experiments. A.F.M. performed the homology modeling and contributed to writing the manuscript. C.J.M. and G.M.B. primarily wrote the manuscript with input from H.Q. and A.F.M.

## Data Availability Statement

All data generated during this study are included in the figures and supplementary figures. Any materials, such as antibodies or nematode strains generated during this study, are available from the corresponding author upon request.

