## Supplemental Figures for "UNC-45 has a crucial role in maintaining muscle sarcomeres during aging in *Caenorhabditis elegans*"

### Table S1. Primers

| gene | Forward primer | Reverse primer |
|---|---|---|
| *gapdh-2* | 5' ACTCTTCACGATTGTCGCTTAGT 3' | 5' AACCCACATACGGACGGTTG 3' |
| *ges-1* | 5' GCTAAAACCGGAGTTCCCCAA 3' | 5' CGTTCCAGAAAGCGAGAGGT 3' |
| *hsp-90* | 5' ATTCGCTACCAGGCACTCAC 3' | 5' GACAAGATCGGCCTTGGTCA 3' |
| *unc-45* | 5' TGCTGAAGGTGGAACGGTTT 3' | 5' CGTATGCTCGTTGTCCAGGG 3' |
| *unc-54* | 5' ATCCAAGCCATTCATGCCGA 3' | 5' GGTCAACGTGCTGAGAGTGT 3' |
| *myo-3* | 5' TCCAGAAGATGGATTCGTCGC 3' | 5' GCGGGTTCATCTCTTGGCAT 3' |

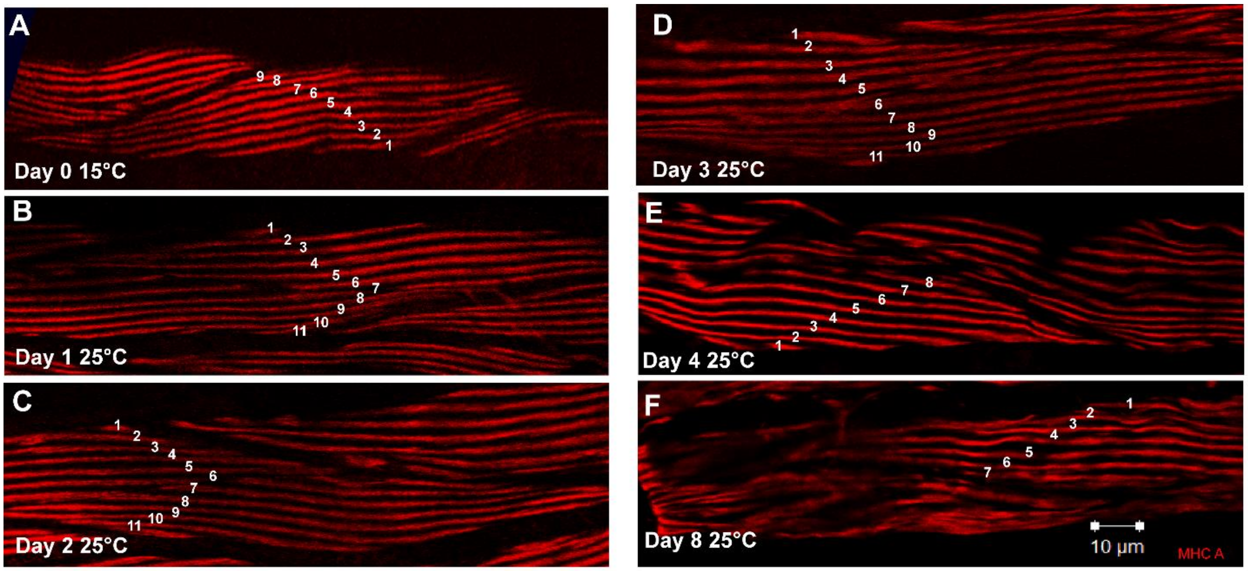


**Figure S1. *unc-45(e286)* representative images of MHC A immunostaining to count A-bands as an assessment of assembled thick filament quantity.** A-F) are representative images of body wall muscle near the vulva immunostained with anti-MHC A at different ages of adulthood (day 0, 1, 2, 3, 4, 8) and grown at either 15ºC (A) or 25ºC (B-F) with an A-band count depicted as white numbers along the A-bands.

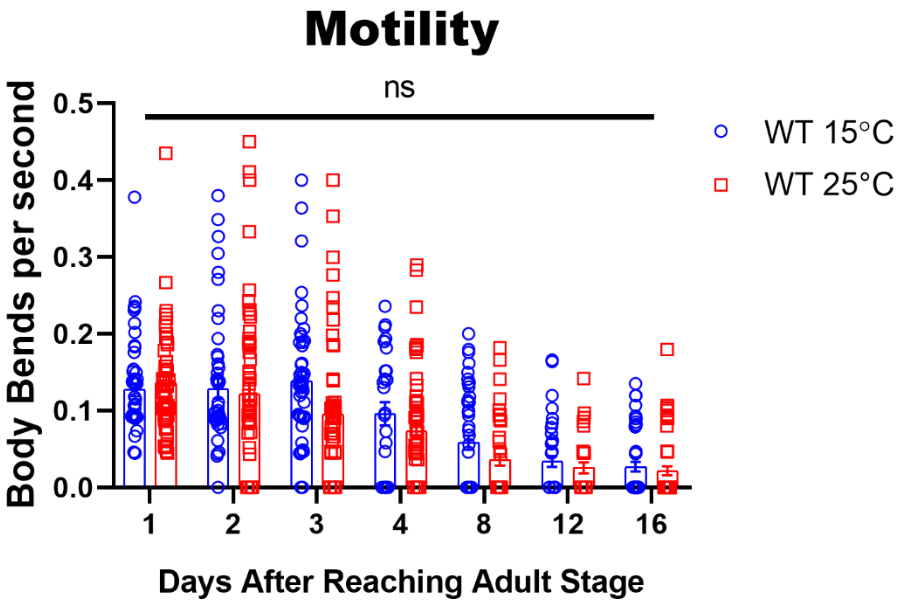


**Figure S2. Crawling motility does not change significantly when wildtype worms are grown at 15°C vs. 25°C.** Quantification of agar crawling motility assays at different ages of adulthood and grown at either 15ºC or 25ºC measured in body bends per second.

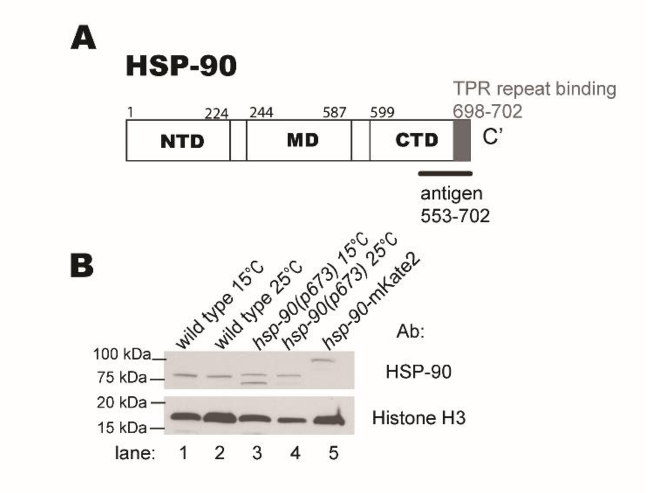


**Figure S3. HSP-90 Antibody validation.** A) shows the protein domains of HSP-90 and the epitope that was used as an antigen when creating the antibody. B) shows a Western Blot validating the antibody that was produced. shows HSP-90 loss of function temperature sensitive mutants (lanes 3-4) compared to wild type (lanes1-2) to show that the mutants have altered HSP-90 and one CRISPR generated fluorescently tagged (mKate2) HSP-90 strain (lane 5) compared to wild type (lanes 1-2) to show that the antibody recognizes one protein band of the appropriate size (106.3 and 80.3 kDa respectively).

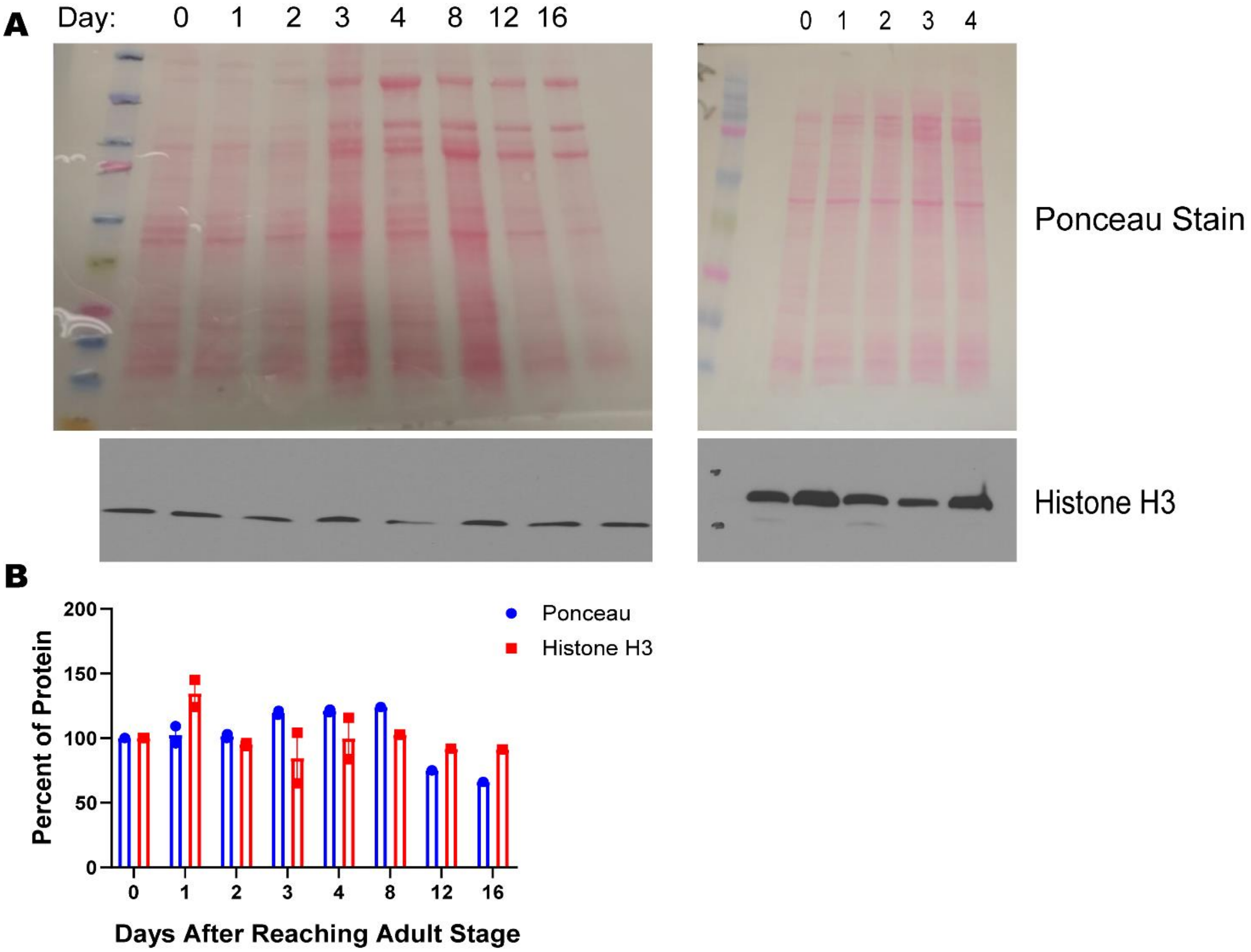


**Figure S4. Histone H3 compared to Ponceau S staining.** A) depicts Ponceau S staining above the Histone H3 from the same blot. B) depicts quantification of Ponceau S staining levels compared to Histone H3 levels at different ages. Fiji Image J was used to measure protein levels.

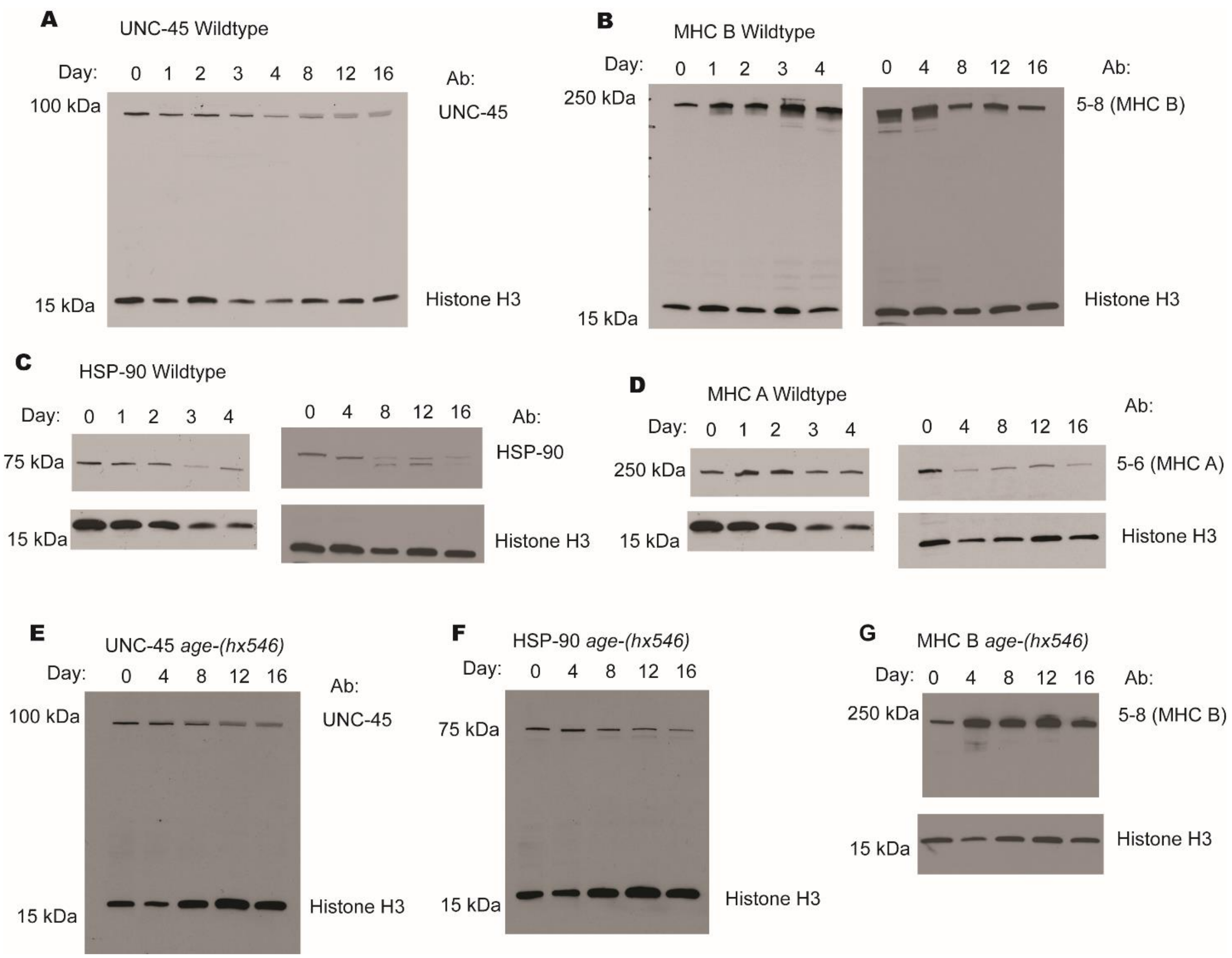


**Figure S5. Representative Western Blot images from wildtype and *age-1(hx546)*.** A-D) depict representative images of UNC-45 (A), MHC B (B), HSP-90 (C), and MHC A (D) Western Blots from wildtype worms at different ages. E-G) depict representative images of UNC-45 (E), HSP-90 (F), and MHC B (G) from *age-1(hx546)* worms at different ages. The load control Histone H3 is also shown for each blot. When blots are shown as two images instead of one, it is because a different exposure time was needed for the two different proteins on the blot.

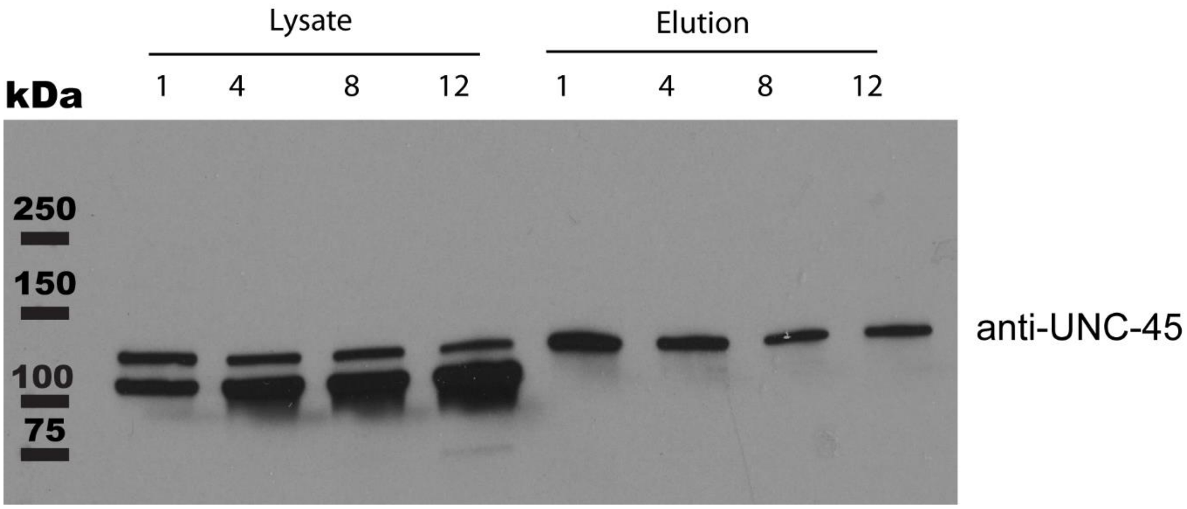


**Figure S6. mNeonGreen Immunoprecipitation.** Part of the protein lysate (~1%) of each sample used for the immunoprecipitation (right side of blot) and part of the total elution (~12.5%) from each immunoprecipitation (left side of blot) were run on a 4-20% SDS-PAGE gel, transferred to nitrocellulose, and reacted with the UNC-45 antibody to ensure that UNC-45::mNeonGreen was pulled down with the pre-conjugated mNeonGreen nanobodies magnetic beads.

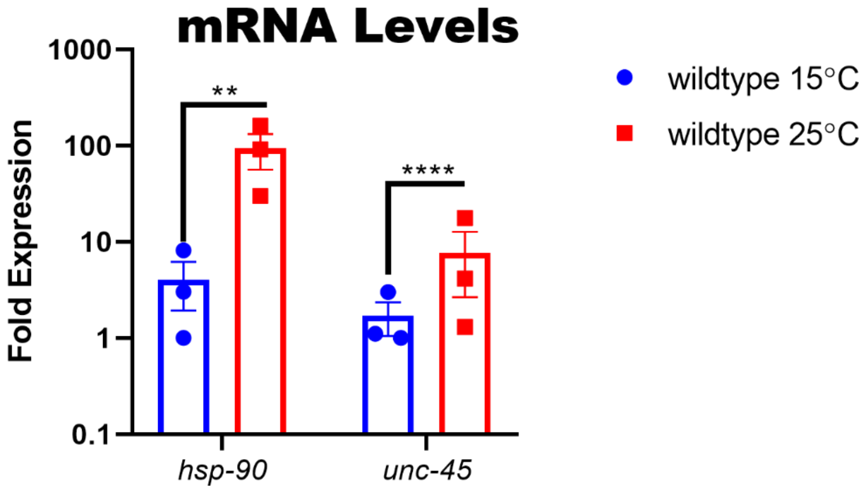


**Figure S7. Steady state transcript expression of *hsp-90* and *unc-45* in Wildtype worms grown at 15ºC and 25ºC.**
